# Parkinson’s disease modeling in regenerative spiny mice (*Acomys dimidiatus*) captures key disease-relevant behavioral, histological, and molecular signatures

**DOI:** 10.1101/2025.11.06.687049

**Authors:** Sayan Dutta, Marion Pang, Renée R. Donahue, Tsui-Fen Chou, Ashley W. Seifert, Viviana Gradinaru

**Affiliations:** Division of Biology and Bioengineering, California Institute of Technology, Pasadena, CA, 91125, USA; Aligning Science Across Parkinson’s (ASAP) Collaborative Research Network, Chevy Chase, MD 20815; Department of Biology, University of Kentucky, Lexington, KY 40508, USA

## Abstract

Parkinson’s disease (PD) is a multifactorial neurodegenerative disorder that has been modeled extensively in animals, primarily rodents, but also in non-human primates and non-mammalian organisms. However, no single animal model fully recapitulates the hallmarks of PD pathology. Here, we extend this work by modeling PD for the first time in the spiny mouse (*Acomys dimidiatus*), a mammal notable for its robust regeneration of multiple tissues. We show that the nigrostriatal pathway of *A. dimidiatus* is vulnerable to both acute 6-hydroxydopamine (6-OHDA) toxicity and chronic α-synuclein (αSyn) preformed fibril-induced aggregation. Mouse αSyn PFFs produced widespread pS129-positive αSyn inclusions across multiple brain regions, mirroring a key pathological hallmark of PD. Compared to C57BL/6J mice, *A. dimidiatus* exhibited more pronounced behavioral impairments, greater nigrostriatal degeneration, and higher pS129-αSyn inclusion burden within substantia nigra pars compacta (SNpc) neurons. To probe the molecular underpinnings behind the vulnerability, we performed single-cell spatial proteomics, which revealed extensive proteomic alterations in dopaminergic neurons associated with αSyn aggregation. Multiple proteins were dysregulated in *A. dimidiatus*, including those involved in proteasomal function, mitochondrial pathways, and oxidative stress regulation, which are processes commonly implicated in PD. Notably, proteomic analysis identified heightened astrocytic activation in the SNpc, which we validated histologically, suggesting a distinct glial response compared to mice. Together, these findings expand our understanding of PD-relevant pathophysiology across species and establish *A. dimidiatus* as a model for studying mechanisms of neurodegeneration.

## Introduction

Parkinson’s disease (PD) is a progressive neurodegenerative disorder primarily characterized by the selective degeneration of dopaminergic neurons in the substantia nigra pars compacta (SNpc) and the pathological accumulation of misfolded α-synuclein (αSyn) in the form of Lewy bodies and neurites^1,2^. Over the past few decades, preclinical animal models, especially in rodents, have proven indispensable in elucidating mechanistic insights^3,4^ and testing therapeutic interventions^5,6^. Toxin-induced models such as 6-Hydroxydopamine (6-OHDA) and 1-Methyl-4-phenyl-1,2,3,6-tetrahydropyridine (MPTP) have provided key insights into dopaminergic vulnerability and oxidative damage^7^, while αSyn-based models using viral vectors or preformed fibrils (PFFs) have allowed the study of proteinopathy and propagation^8^.

In addition to rodents, other model organisms such as non-human primates, zebrafish, and *Caenorhabditis elegans* have provided unique contributions to PD research. Non-human primate models have been instrumental by recapitulating motor symptoms and cortical pathology with high translational fidelity, aiding the development of deep-brain stimulation and dopaminergic therapies^9,10^. Zebrafish, due to their optical transparency and conserved dopaminergic circuitry, have facilitated high-throughput screening^11^. *C. elegans* have allowed the discovery of genetic modifiers of αSyn toxicity and mitochondrial pathways due to their genetic tractability and simplicity^12^. Each species offers distinct advantages that have complemented rodent studies and enriched our understanding of PD at cellular, circuit, and behavioural levels.

Our current work aims to expand the preclinical modeling toolbox for PD by introducing a novel species: the spiny mouse (*Acomys dimidiatus*). *Acomys* are unique among adult mammals in their exceptional regenerative capacity, including scar-free wound healing and complex tissue regeneration^13,14^ and regrowth of axonal fibers following spinal cord injury^15^. These regenerative traits are accompanied by distinct mitochondrial and cellular stress responses and inflammation^16^, which are key processes implicated in neurodegenerative disease mechanisms^17^. Therefore, *Acomys* might offer a unique opportunity to explore both fundamental disease pathways and potential regenerative strategies for the central nervous system (CNS) circuits implicated in PD and other neurodegenerative disorders.

As a first step in this direction, we modeled PD in *A. dimidiatus* using 6-OHDA or αSyn-PFFs, successfully recapitulating key disease hallmarks including behavioral deficits, nigrostriatal degeneration, and αSyn aggregation. To contextualize these findings, we performed one of the first direct interspecies comparisons in PD, between *A. dimidiatus* and the widely used C57BL/6J laboratory strain of *Mus musculus*. Finally, using single-cell spatial proteomics^18^, we identified molecular alterations in pathology-enriched regions compared to non-pathological tissue, identifying potential disease-relevant pathways in a spiny mouse model of PD.

## Results

### Spiny mice *(Acomys dimidiatus)* exhibit nigrostriatal vulnerability to both toxin- and proteinopathy-based PD stressors

To evaluate the susceptibility of the *A. dimidiatus* (Fig. 1A) nigrostriatal pathway to representative Parkinsonian insults, animals were stereotaxically injected unilaterally in the striatum with either 6-OHDA or αSyn PFFs, with the contralateral hemisphere serving as an internal control (Fig. 1B). Dosing was estimated from studies with established rodent models^19,20^, and targeting was guided by conserved anatomical landmarks^21^.

**Figure 1.**
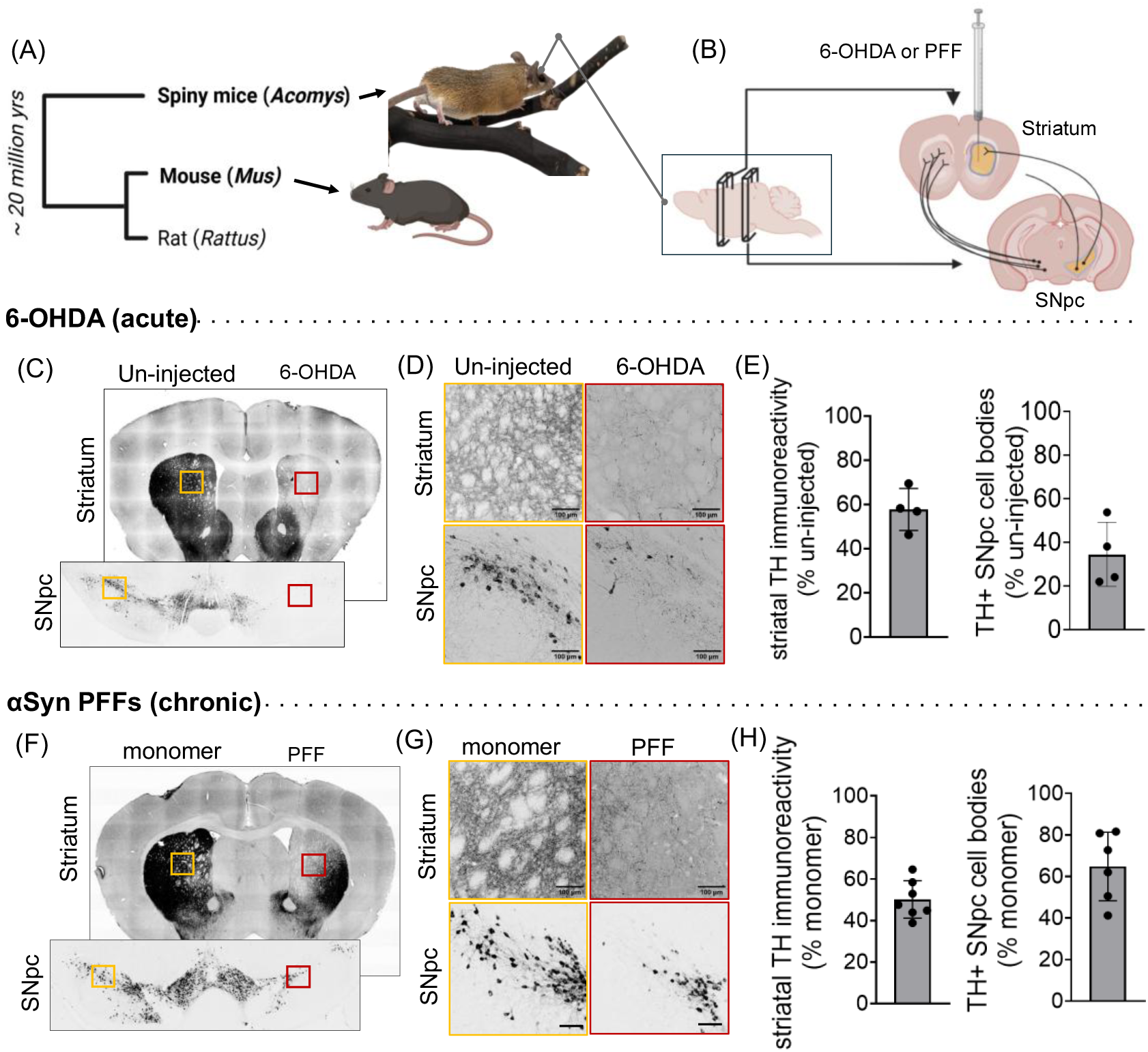
The spiny mouse nigrostriatal pathway degenerates following exposure to 6-OHDA or recombinant mouse αSyn PFFs: (A) Representative image of spiny mice (*Acomys*) and a simplified phylogenetic tree depicting the evolutionary divergence from commonly used laboratory rodents (*Mus* and *Rattus*). (B) Schematic of coronal brain sections used to assess nigrostriatal integrity. The striatal section shows the injection site for 6-OHDA or αSyn PFFs, and the midbrain section highlights the SNpc containing dopaminergic cell bodies projecting to the striatum. Left, intact nigrostriatal circuit; right, lesioned pathway illustrating degeneration. (C) Representative anti-TH immunohistochemical (IHC) staining showing loss of dopaminergic terminals in the striatum and cell bodies in the SNpc of the 6-OHDA-injected hemisphere. (D) Higher-magnification views highlighting TH loss in the regions boxed in (C). Scale bars: 100 μm. (E) Quantification of TH-positive terminal loss in the striatum and dopaminergic cell-body loss in the SNpc following 6-OHDA injection (n=4 animals). (F) Representative anti-TH IHC showing reduced striatal TH density and SNpc dopaminergic neuron loss in the PFF-injected hemisphere versus the monomer-injected control side 6 months post-injection. (G) Higher-magnification views highlight degeneration in the regions boxed in (F). Scale bars: 100 μm. (H) Quantification of striatal TH-terminal density and SNpc cell-body counts following PFF injection (n=6-7 animals). Schematics were prepared using BioRender

Intrastriatal injections of 12µg 6-OHDA induced nigrostriatal degeneration, marked by loss of tyrosine hydroxylase (TH) immunoreactivity in the striatum and reduced dopaminergic neuron counts in the SNpc 4 weeks post-injection (Fig. 1, C-D). Quantification revealed approximately 40% loss of striatal TH density and ∼60% loss of TH^+^ cell bodies in the SNpc (Fig. 1E). Overall, these observations are consistent with 6-OHDA models in *Mus*^22^ and *Rattus*^23^, indicating sensitivity to oxidative dopaminergic injury.

Next, we assessed the response of the *Acomys* nigrostriatal pathway to a slower, proteinopathy-based PD model using recombinant mouse αSyn PFFs ^8,19,20^. Because *A. dimidiatus* αSyn is identical in sequence to *M. musculus* αSyn, we used mouse PFFs, which are widely adopted for modeling synucleinopathy^24,25^. This model exploits the seeding capability of exogenous fibrils to induce misfolding and aggregation of endogenous αSyn, ultimately leading to gradual neurodegeneration. *A. dimidiatus* received bilateral intrastriatal injections, with PFFs in one hemisphere and an equivalent amount of monomeric αSyn in the contralateral side as control^4^. At 6 months post-injection (m.p.i.), a time point with detectable nigrostriatal degeneration established in murine studies^26^,we observed a significant reduction in striatal TH fiber density and corresponding loss of SNpc neurons, restricted to the PFF-injected hemisphere (Fig. 1, F-G). Quantification revealed ∼50% loss of striatal TH density and ∼40% loss of TH^+^ cell bodies in the SNpc (Fig. 1H).

Together, these findings establish that *Acomys* SNpc neurons are susceptible to both acute oxidative stress-driven and chronic αSyn aggregation-driven degeneration.

### Intrastriatal PFF injection triggers phosphorylated αSyn inclusions throughout the brain

Along with nigrostriatal degeneration, PFFs also generated endogenous αSyn inclusions immunopositive for serine-129–phosphorylation (pS129-αSyn), a pathological hallmark of PD, as previously demonstrated in other rodent^19,26^ and non-human primate models^10^. We tested both one-site and two-site unilateral intrastriatal injections—the latter strategy often used in larger animal models to enhance seeding efficiency ^19,27,28^ (Supp. Fig. 2). pS129-αSyn inclusions were detectable as early as 1 m.p.i.—the earliest time point assessed—and persisted up to at least 6 m.p.i., consistent with murine PFF models^26,27^ (Fig. 2, A-B). pS129-αSyn inclusions were identified throughout multiple brain regions, including sensorimotor cortex, amygdala, striatum, and SNpc (Fig. 2A). At 1 m.p.i., inclusions in some brain regions (e.g., cortex, striatum) were predominantly localized to neurites, but by 6 m.p.i. they were also evident in somata(Fig. 2B). Depending on the PFF dose, as many as 40–90% of TH⁺ dopaminergic neurons in the *A. dimidiatus* SNpc developed pS129-αSyn somatic inclusions (Supp. Fig. 2). In contrast to other regions, the SNpc showed a decline in detectable aggregates between 1 and 6 m.p.i., likely reflecting degeneration and loss of αSyn-laden dopaminergic neurons, as previously reported in rodent studies^26,27^.

**Figure 2.**
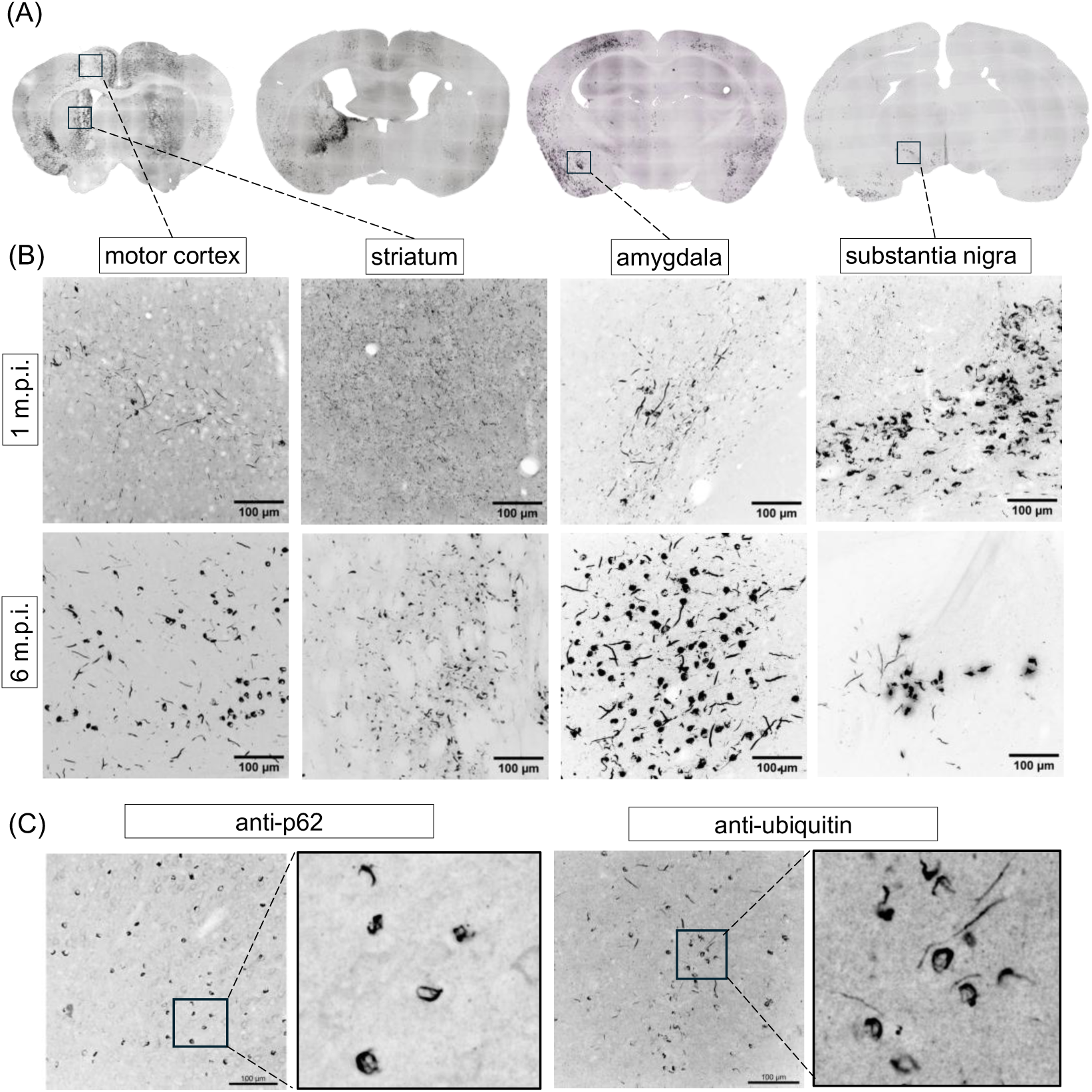
Striatal PFF injection in *A. dimidiatus* induces PD-like pathology across brain regions. (A) Representative coronal brain sections showing pS129-αSyn immunoreactivity in the cortex, striatum, amygdala, and substantia nigra after αSyn PFF injection into the striatum. (B) Representative (from n=3 animals per timepoint) higher-magnification images of corresponding brain regions at 1- and 6-months post injection (m.p.i.), illustrating progressive accumulation of pS129-αSyn. Scale bars: 100 µm. (C) Representative (from n = 3 animals) images showing accumulation of p62- and ubiquitin-positive inclusions in the cortex at 6 m.p.i.

In addition to pS129-αSyn, the aggregates displayed several hallmark features of Lewy-like pathology. Immunohistology revealed polyubiquitination and accumulation of the autophagy adaptor p62 (Fig. 2C), closely mirroring pathological signatures observed in other synucleinopathy models^29,30^ and in human PD tissue^31^. Together, these results confirm that spiny mice can effectively recapitulate the key molecular and anatomical hallmarks of PD-associated pathology.

### PFF-treated *A. dimidiatus* exhibit stronger motor behavioral deficits than C57BL/6J mice

We next assessed motor behavioral deficits, one of the hallmarks of PD, in the PFF model. For comparison, we also assessed the commonly used C57BL/6J mouse strain. 7.5 µg of αSyn PFF was injected unilaterally in both *A. dimidiatus* and C57 animals (with the contralateral side receiving monomeric αSyn as a control) to induce a nigrostriatal lesion in one hemisphere. Motor behavior was assessed at 1 m.p.i., an early timepoint when motor deficits are typically minimal^19,20,24^, and again at 6 m.p.i. (Fig. 3A).

**Figure 3:**
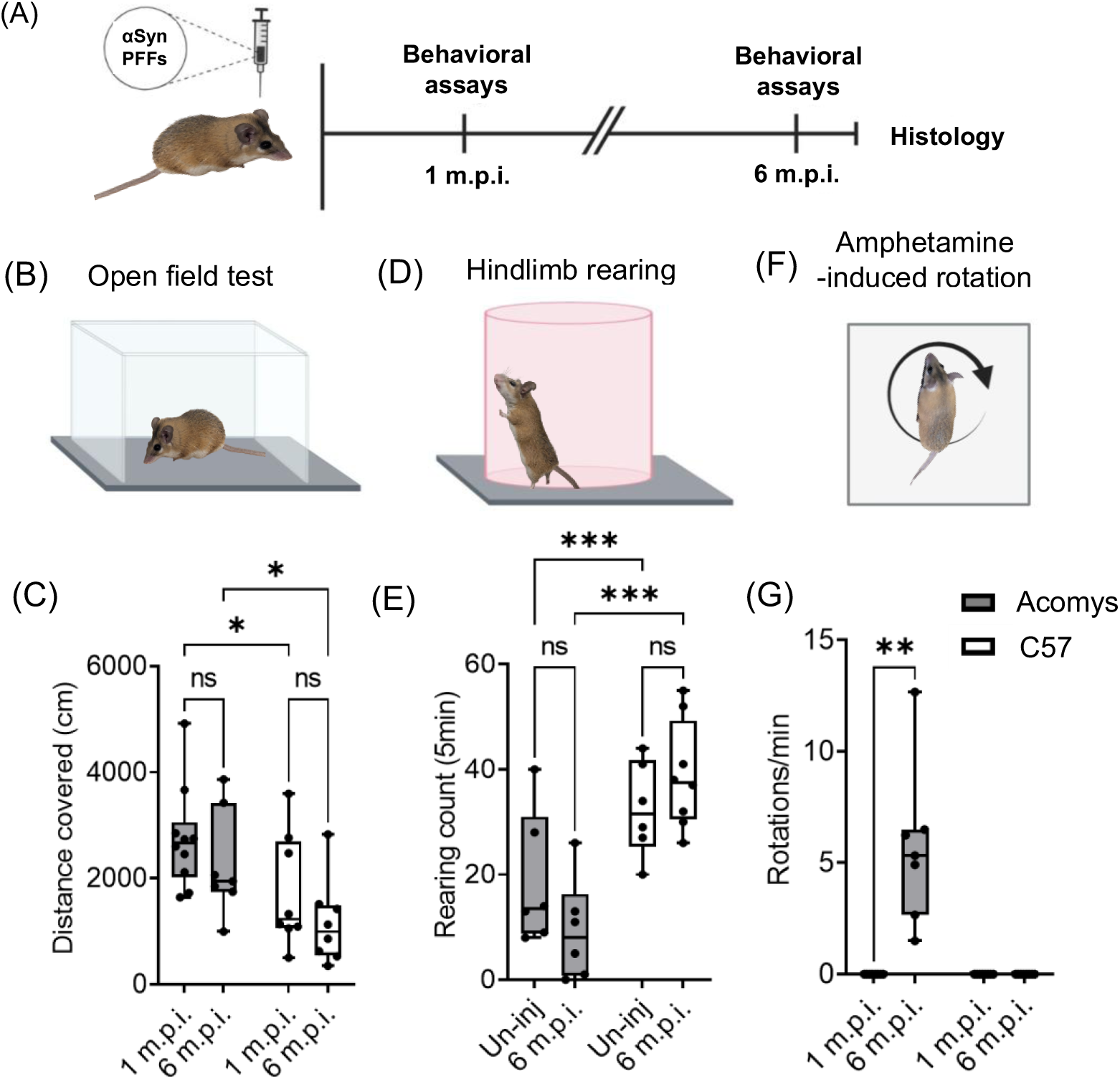
Motor behavioral assessment of *A. dimidiatus* and *M. musculus* following unilateral PFF injection: (A) Schematic of timeline of behavioral experiments conducted in spiny mice and C57BL/6J mice over the post-injection period. (B) Illustration of the open field test setup and (C) quantification of total distance traveled in the open field arena at 1- and 6- m.p.i. (D) Diagram depicting spontaneous vertical rearing behavior test (cylinder test) and (E) rearing event counts in injected animals at 6 m.p.i. (F) Schematic of the amphetamine-induced rotation assay and (G) quantification of rotations per minute in the ipsilateral direction relative to the PFF-injected side. Statistical significance was assessed using Student’s *t*-test (*p < 0.05; **p < 0.01; ***p < 0.001). Illustrations and schematics were prepared using BioRender.

First, we evaluated gross motor function using the open field test (Fig. 3B). Neither species showed significant differences in locomotor activity between 1 m.p.i. and 6 m.p.i. measurements. Nevertheless, clear baseline species differences were evident, with *A. dimidiatus* exhibiting greater locomotor activity than C57 mice (Fig. 3C). In the rearing test, we observed a modest, nonsignificant reduction in rearing for injected *Acomys* at 6 m.p.i compared to uninjected animals (Fig. 3, D-E). Again, task-dependent species differences were apparent, with lower baseline rearing counts in *A. dimidiatus* than C57 mice.

Both baseline locomotion and rearing behavior are governed by activity from both sides of the brain. Therefore, for a hemisphere-specific assay, we performed the amphetamine-induced rotation test (Fig. 3F)^32^ to assess the extent of degeneration in the PFF-injected hemisphere compared to the intact side. Strikingly, at 6 m.p.i., *A. dimidiatus* exhibited significant ipsilateral rotation, which has been previously established in other models to indicate at least 40-50% striatal terminal loss compared to the control hemisphere^8,32^ (Fig. 3G). Under identical conditions, C57 mice did not show any rotational behavior.

These results indicate that the PFF model of PD can induce motor behavioral deficits in *Acomys*, and that these deficits may in fact be more severe than in *Mus*. Moreover, the baseline behavioral differences we observed underscore the importance of caution in comparing outcomes across species.

### PFFs induce greater striatal degeneration and pS129-αSyn inclusions in *A. dimidiatus* than C57 mice

To gain histological insight into the interspecies difference we observed in the amphetamine rotation assay, we collected brains after the 6 m.p.i. behavioral assessments. Nigrostriatal degeneration was evaluated by quantifying the reduction in TH immunoreactivity in the striatum, a measure that correlates well with the severity of amphetamine-induced rotation^32^. In both *A. dimidiatus* and C57 mice, the PFF-injected hemisphere showed a marked reduction in TH immunoreactivity relative to the control hemisphere (Fig. 4, A-B); however, *A. dimidiatus* exhibited significantly greater terminal loss than C57 mice (Fig. 4B), despite comparable baseline TH density in the control side of the brain between the species (Fig. 4C).

**Figure 4:**
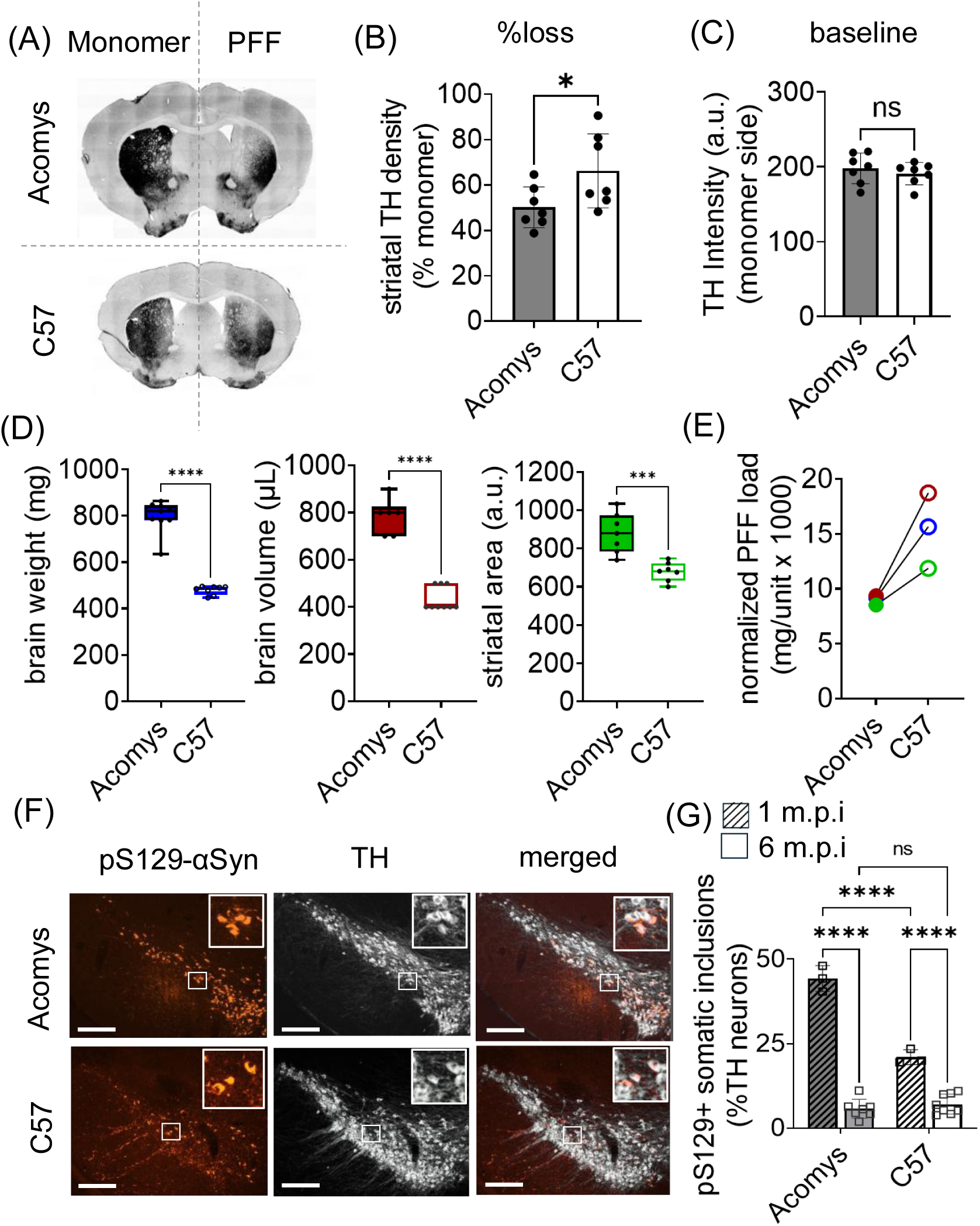
Histological comparison of striatal TH loss and pS129-αSyn pathology between *A. dimidiatus* and C57 mice supports interspecies difference: (A) Striatal TH immunoreactivity at 6 m.p.i., comparing PFF-injected and control (monomer-injected) hemispheres. (B) Quantification of striatal TH density on the injected side, normalized to the intact hemisphere. (C) Striatal TH density in the control side. (D) Baseline comparison of brain weight, brain volume (estimated by water displacement; 100 μL minimum measurable unit), and cross-sectional area of the striatal region at the injection site (from coronal sections) between *A. dimidiatus* and C57 mice. (E) Normalized PFF dose used in this study relative to the average values from panel D. (F) Representative immunohistochemistry images of SNpc showing TH and pS129-αSyn staining (inset: magnified view of the boxed region). Scale bars: 200 μm. (G) Quantification of TH+ neurons containing pS129-αSyn aggregates in the SNpc at 1 and 6 months post-PFF injection. Statistical significance was determined by Student’s *t*-test (ns: not significant; *p < 0.05; ***p < 0.001; ****p < 0.0001).

Interestingly, the greater loss of TH immunoreactivity in *A. dimidiatus* resulted from a lower effective PFF dose. Although the same absolute dose of 7.5 µg of PFF was administered to both species, *A. dimidiatus*—having a larger brain (greater weight, volume, and striatal cross-sectional area at the injection site) (Fig. 4D)—received less PFF per unit volume of brain parenchyma than the C57 mice (Fig. 4E). This indicates that *Acomys* develop more severe striatal degeneration from a given PFF burden than *Mus*; in contrast, relatively larger PFF doses are required in rats to achieve degeneration comparable to mice^19,27^.

In PFF models, αSyn aggregation load correlates with loss of striatal density^19,27^, and neurons bearing aggregates are selectively more vulnerable than aggregate-free neurons^26,33^. To investigate whether greater aggregation might underlie the greater degeneration observed in *A. dimidiatus*, we analyzed pS129-αSyn inclusions in SNpc neurons. At 1 m.p.i., spiny mice displayed a significantly higher percentage of SNpc neurons containing pS129-αSyn inclusions than mice (Fig. 4, F-G). By 6 m.p.i., both species showed markedly reduced numbers of SNpc neurons, with no significant difference between them (Fig. 4G). These findings suggest that *A. dimidiatus* lost a greater proportion of aggregate-bearing SNpc neurons over time^26^, resulting in the more pronounced striatal TH density loss we observed in this species.

Together, these findings highlight a possible greater susceptibility of *A. dimidiatus* to αSyn pathology and support its use as a novel model to study mechanisms of neurodegeneration in PD.

### Single-cell spatial proteomics identifies PD-linked proteins and pathways altered in *A. dimidiatus* pS129-αSyn inclusion-bearing neurons

To define molecular changes in this novel *Acomys* PD model, we profiled proteomic alterations in diseased neurons. For this, we applied a single-cell spatial proteomics workflow that we previously established for formalin-fixed CNS tissue^18^ following immunohistochemistry for TH and pS129 αSyn to capture molecular alterations in aggregate-containing (Agg+) neurons post-PFF injection compared to non-aggregate-containing (Agg-) neurons in the control side of the same animal (Fig. 5A). This approach overcomes limitations of bulk proteomics such as insufficient resolution to resolve altered signatures in the small fraction of aggregate-containing neurons within bulk lysates and potentially confounding effects of differences in tissue composition (e.g., neuron–astrocyte ratios) between *Acomys* and *Mus*. In the absence of an *Acomys* reference proteome, protein identification was performed using the *Mus musculus* Uniprot Swiss-Prot reviewed proteome (canonical & isoform) as a surrogate. We chose this approach rather than deriving a custom *Acomys* database from transcriptomic data, since substantial manual curation would be necessary to achieve reliable annotations. As a caveat, however, sequence divergence between *Acomys* and *Mus* likely reduced identification confidence and increased the apparent false discovery rate (FDR), resulting in fewer matched proteins.

**Figure 5.**
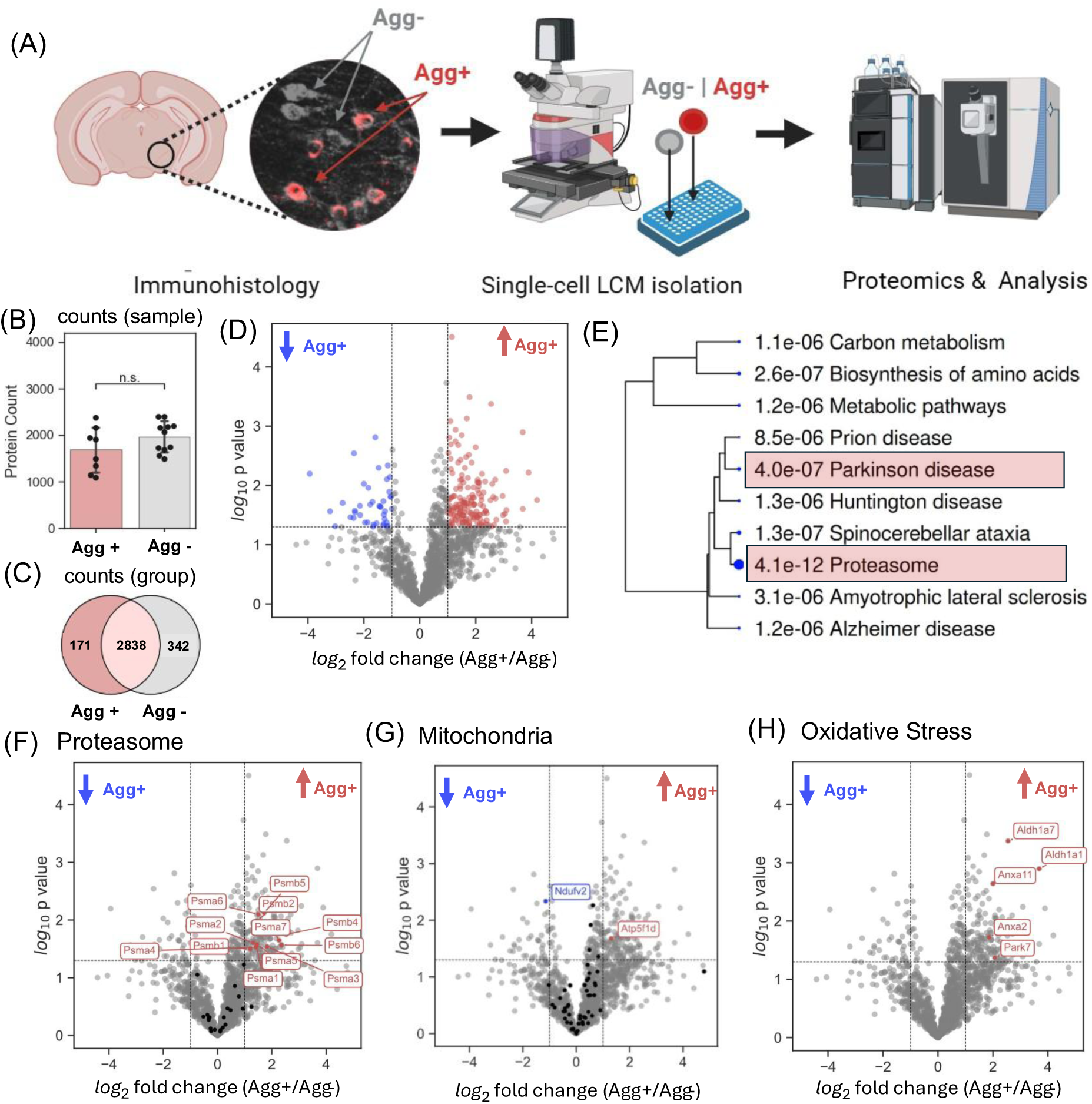
Histology-guided spatial single-cell proteomics reveals dysregulated pathways associated with αSyn aggregation in SNpc neurons of *A. dimidiatus*. (A) Schematic overview of the experimental workflow. Schematic prepared using BioRender. (B) Bar graph indicating the number of proteins per sample identified in aggregate-containing (Agg⁺) and aggregate-free (Agg⁻) neurons, representing mean ± SEM from 9–11 single cells per group. (C) Venn diagram illustrating the overlap of common and unique proteins across all measured cells in the two experimental groups. (D) Volcano plot depicting global protein expression changes, with log₂ fold change on the x-axis and –log₁₀ p-value on the y-axis; upregulated proteins in Agg⁺ neurons are shown in red, downregulated proteins in blue. (E) Hierarchical clustering of significantly altered proteins (p ≤ 0.05, fold change ≥ 2), with the top two most significantly upregulated KEGG pathways highlighted, reveals potentially co-regulated modules. (F–H) Volcano plots highlighting significantly altered proteins associated with key cellular processes, including the proteasome (F), mitochondrial complex (G), proteins linked to managing oxidative stress (H). . Black points indicate other, non-significant protein hits from the same pathways.

We identified on average ∼2,000 unique proteins per single neuron (Fig. 5B) and ∼3,000 protein groups across both neuronal populations, with 342 proteins detected exclusively in Agg-neurons and 171 proteins unique to Agg+ neurons (Fig. 5C; n = 9–11 single neurons per group). After further filtering for proteins detected in > 40% of samples within at least one group, 165 proteins were significantly up-regulated and 41 down-regulated (p < 0.05, log_2_(FC) ≥ 1) in Agg+ neurons relative to Agg-ones (Fig. 5D). Pathway enrichment analysis of the significantly altered hits against KEGG pathways, a curated collection of molecular networks from experimentally validated literature^34^, highlighted “Proteasome” and “Parkinson’s disease” as among the most significantly altered pathways (Fig. 5E). Proteasomal dysfunction, which was reflected by upregulated proteasomal subunits (e.g., PSMB5, PSMA6) in Agg+ neurons (Fig. 5F), also emerged as a key component of the Parkinson’s disease pathway (Supp. Fig. 3). In contrast, proteins from mitochondrial complex-I (Ndufv2) was down-regulated and complex-V (Atp5f1d) was upregulated (Fig. 5G), pointing to possible alteration of mitochondrial function. Upregulation of some oxidative stress managing proteins (Aldh1a1, DJ-1 (PARK7)) was also observed, possibly indicating an increased reactive oxidative species buildup in aggregated-containing neurons.

Because the brain is a highly heterogeneous and interwoven tissue, single-neuron proteomics can capture low-level signals from adjacent cell types. In our previous single-cell C57 mouse study, we detected glial markers in neuron-enriched samples^18^, and we saw a similar pattern here in *A. dimidiatus*. For example, mass-spectrometry revealed greater GFAP abundance in Agg+ *A. dimidiatus* compared with C57 mice, where no significant increase was observed (Fig. 6A). Follow-up immunohistochemistry confirmed reactive astrogliosis, elevated GFAP immunoreactivity in the PFF-injected hemisphere relative to the control hemisphere, in *A. dimidiatus* but not in C57 mice (Fig. 6B). Quantification demonstrated an approximately 50% increase in GFAP intensity, with a visibly higher density of GFAP-positive astrocytes in the substantia nigra (Fig. 6C). In contrast, microglial activation, assessed by IBA1 immunoreactivity, did not show a significant elevation in either species at the 6-month time point (Fig. 6D). C57BL/6J mice exhibited no change in reactive microglial profile, consistent with a recent study^24^, whereas *A. dimidiatus* showed only a subtle, non-significant increase in ramified microglia on the PFF-injected side (Supp. Fig. 4).

**Figure 6.**
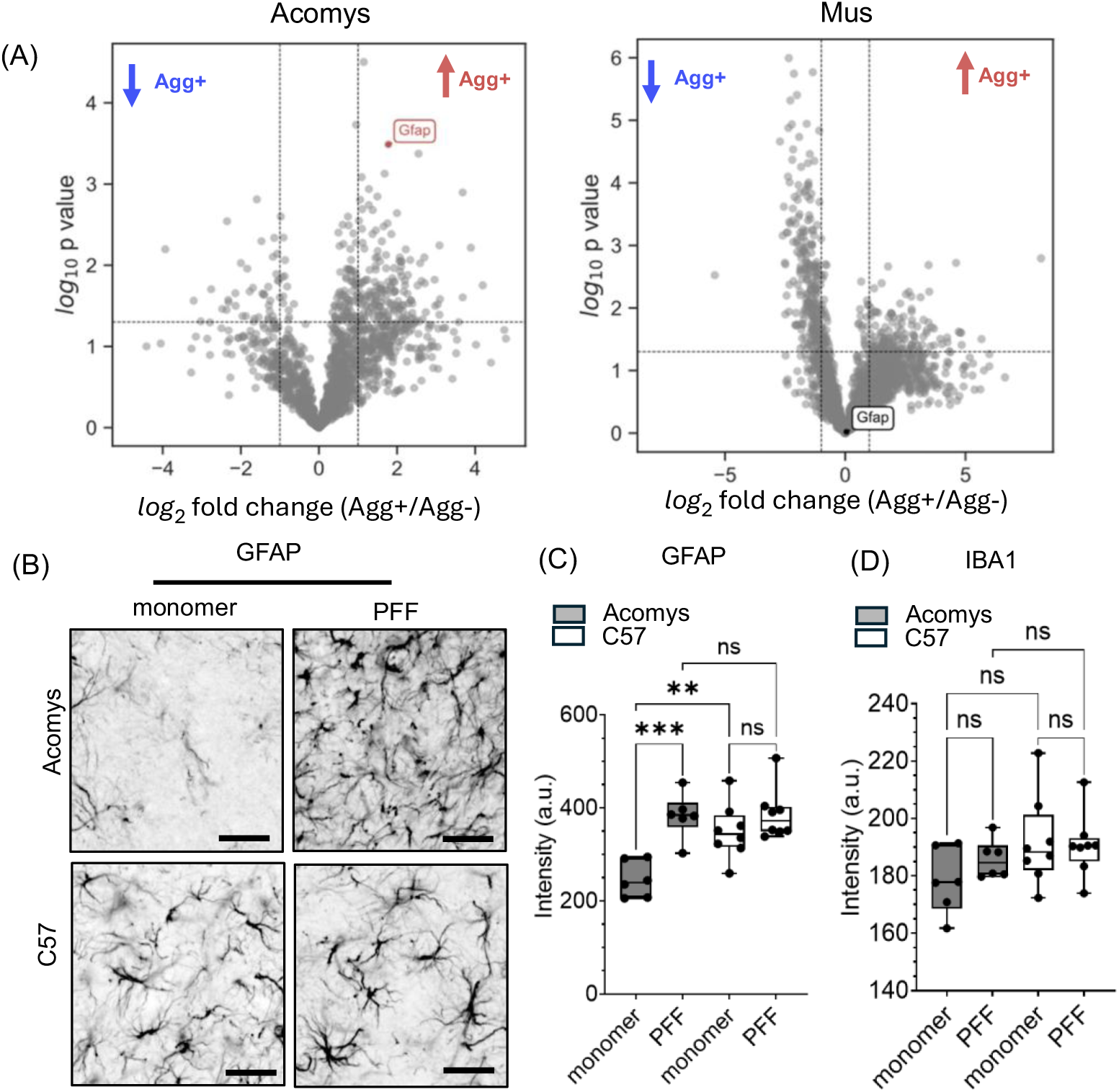
Glial alterations in *A. dimidiatus* and C57BL/6J mice at 6 months post-PFF injection. (A) Volcano plot showing significant fold-change alteration in glial fibrillary acidic protein (GFAP). (B) Representative GFAP immunostaining in the substantia nigra of spiny mice and C57BL/6J mice (scale bars, 25 µm). (C) Quantification of GFAP signal intensity in the PFF-injected region in both species. (D) Quantification of IBA1 signal intensity in the substantia nigra of spiny mice and C57BL/6J mice.

In line with the concept of multi-pathway failure in PD^35^, these findings reveal a coordinated disruption of proteasomal, mitochondrial, oxidative pathways and altered immune response, core processes implicated in PD pathogenesis, in aggregate-bearing samples of PFF-treated *A. dimidiatus*.

## Discussion

Spiny mice (*Acomys* spp.) are increasingly recognized as a powerful emerging mammalian model due to their exceptional regenerative capacity across multiple organs^15,36–38^, including post-injury regeneration of skin, kidney, and liver^14,39–41^, and partial functional recovery after spinal cord injury^15^. Building on this foundation, we undertook the first systematic assessment of whether *Acomys* can model PD-like pathology, especially neurodegeneration, extending its use to CNS disorders. We found that *A. dimidiatus* treated with either 6-OHDA or αSyn PFFs developed pronounced PD-like features, including robust nigrostriatal degeneration (Fig. 1) and intraneuronal proteinaceous aggregates (Fig. 2), grossly resembling the phenotypes reported in comparable mouse and rat models^7,19,20,28^. Within the timeframe examined (6 months), however, we did not observe clear evidence of neuronal protection or regeneration—consistent with the limited capacity of post-mitotic neurons to rebuild circuits once lost. This suggests that the chronic, intracellular nature of αSyn PFF or 6-OHDA insults differs fundamentally from the acute localized axonal damage in spinal cord injury models, where regeneration can be driven by intact cell bodies^42,43^. These observations establish that, despite the remarkable regenerative potential of its peripheral tissues, *A. dimidiatus* does not display overt neuroprotection or neuroregeneration in a PD-like setting.

In line with recent comparative studies in spiny mice^15,44^, we systematically evaluated how *A. dimidiatus* compares with one of the wild-type murine strains most widely used for modeling PD, C57BL/6J *M. musculus*. Assessing behavior, approximately 6 months following PFF injection, neither species showed significant deficits in spontaneous movement tasks, including the open field and cylinder tests (Fig. 3C, 3E), in agreement with previous rodent studies ^19,20,27,45^. A recent mouse study^24^ using a higher PFF dose (with correspondingly greater striatal TH density loss) did report a decrease in rearing in the cylinder test, underscoring the dose dependence of behavioral outcomes. Interestingly, we noted a similar, though not significant, downward trend in *A. dimidiatus* at our lower dose (Fig. 3E), suggesting greater degeneration relative to *M. musculus*. This became clearer in the induced-movement assay of amphetamine rotation. Here, *A. dimidiatus* exhibited a significant increase in pathological behavior following unilateral PFF injection, with no increase in C57 mice given the same dose, indicating a greater lesion (Fig. 3G). This was corroborated histologically by more extensive striatal TH^+^ nerve terminal loss in *A. dimidiatus* than in C57 mice (Fig. 4, A-B). The complete absence of amphetamine rotation in the C57 mice (Fig. 3G) cohort was initially unexpected, given widespread pathology (Fig. 4, F-G) and significant striatal TH loss (Fig. 4B). However, literature from established unilateral 6-OHDA^32^ and AAV αSyn overexpression models^8^ indicates that robust rotational behavior typically appears only when striatal TH^+^ terminal loss exceeds a threshold of ∼40–50%, which at the dose and six-month time point used here, most *A. dimidiatus*, but not most C57 mice, appeared to cross (Fig. 4B). Nonetheless, we did not observe rotation in individual C57 mice that exhibited comparable striatal TH loss (> 40%) to individual *A. dimidiatus* that did show robust rotation (Fig. 4B), suggesting potential species-specific differences (e.g., processing kinetics) in amphetamine response. Taken together, our amphetamine-induced rotation data combined with the greater degree of striatal TH loss— particularly given lower relative PFF dose in the relatively larger *A. dimidiatus* (Fig. 4E)—point toward more pronounced nigrostriatal degeneration in *Acomys* than in mice under these conditions. Further studies using higher PFF loads, additional PD models, and comparisons with other rodents (including rats) will be necessary to determine whether spiny mice indeed possess a more vulnerable nigrostriatal circuit.

Understanding the mechanism underlying the apparent heightened vulnerability of *A. dimidiatus* to αSyn PFFs requires further exploration. Our data point to a greater burden of αSyn aggregation in *A. dimidiatus* SNpc neurons as one likely contributor (Fig. 4F–G). However, several additional factors could plausibly account for the increased susceptibility, including differences in neuronal uptake of fibrils^46^, variations in endo-lysosomal or protein-clearance pathways^47,48^, and species-specific features of nigrostriatal circuit organization such as dopaminergic cell subtypes or patterns of axonal arborization^49,50^.

To reveal molecular-level differences that might underlie the distinct behavioral and pathological profiles observed, we performed single-cell spatial proteomics^51,52^ on fixed tissues, guided by immunohistochemical staining for TH and pS129 αSyn. This single-cell approach overcomes undersampling issues for the sparse population of aggregate-positive SNpc neurons in bulk approaches. To our knowledge, this represents one of the first applications of single-cell spatial proteomics in a PD model, and the first for *Acomys*. The resulting data revealed striking alterations in proteasomal and mitochondrial proteins following PFF injection, suggesting potential dysfunction in these pathways. αSyn aggregates are known to impair proteasome activity^53^, and PD patients show reduced proteasomal function^54^. Interestingly, in our dataset, we observed increased expression of proteasomal subunits in PFF-treated *A. dimidiatus* (Fig. 5F), which may reflect a compensatory cellular response to aggregation-induced inhibition or an attempt to enhance clearance of misfolded proteins^55^. Likewise, αSyn aggregates are reported to interact with mitochondria^56^, and mitochondrial impairment is a well-established feature of PD^57^; the reduced abundance of mitochondrial complex-I subunits we observed is consistent with this idea.

The glial activation data were particularly noteworthy: while in mice GFAP and IBA1 were not elevated, as in prior unilateral PFF injection studies^24^, in contrast, *A. dimidiatus* mounted a distinct immune response characterized by limited microglial activation but robust astrogliosis. This pattern may be a consequence of greater neuronal degeneration, representing a compensatory attempt to enhance debris clearance^42^, or an inherently different microglial reaction to injury in *Acomys*^15^. Together, these findings underscore the potential of *Acomys* to yield new insights into astrocyte involvement in PD pathogenesis and glia–neuron interactions in synucleinopathy.

Even though *Acomys* did not exhibit overt neuroprotection to PD-relevant insults, these results establish an essential baseline for refining future study designs and expectations for neuroprotection and CNS regeneration work in *Acomys*. Some of our findings, such as extracellular matrix gene upregulation and astrogliosis, underscore the need to investigate extra-neuronal mechanisms that may shape regenerative responses in *Acomys* across diverse types of brain injury. This work also marks an early step in establishing *Acomys* as a model for CNS-targeted investigations, presenting what we believe is the first detailed investigation of brain circuit alterations in response to a PD-relevant insult. Several challenges remain. We found it difficult to target smaller brain structures, such as the medial forebrain bundle or SNpc, based on the currently available stereotaxic atlas^21^, indicating further refinement might be necessary. Secondly, the significant baseline differences we observed between *M. musculus* and *A. dimidiatus* show that further studies are needed to better characterize *Acomys* behavior in comparison with commonly used laboratory models. In general, developing well-curated, species-specific databases will enable greater systems-level insight into CNS regeneration and neurodegeneration in *Acomys*.

## Experimental Methods

### Animal Husbandry

A breeding colony of spiny mice was established at Caltech using breeding pairs transferred from Dr. Ashley Seifert’s laboratory at the University of Kentucky. Animals were housed in a temperature (71-79°F) and humidity (30-70%)-controlled facility under a standard 13/11-hour light/dark cycle. Cage bedding was changed every two weeks to minimize stress while maintaining a stable environment. Non-breeding spiny mice were group-housed in ventilated cages (3–5 animals per cage, same sex) containing nesting materials and environmental enrichment (e.g., wood blocks, shelters, twigs, manzanita wood branches), with ad libitum access to standard rodent chow (Lab Diet #5053) and water (Sup. Fig. 1). All husbandry and experimental procedures were conducted in compliance with the guidelines of the Caltech Office of Laboratory Animal Resources (OLAR) and were approved and monitored by the Institutional Animal Care and Use Committee (IACUC) at Caltech. For interspecies comparative studies, C57BL/6J mice (The Jackson Laboratory #000664, RRID: IMSR_JAX:000664) were housed 4–5 per cage under a 12-hour light/dark cycle. Bedding and cages for C57 mice were changed weekly, and animals were provided with unrestricted access to food and water. Sexually mature 3–6-month-old C57 mice were compared with 6–12-month-old spiny mice, taking into consideration species-specific development and lifespan differences.

### 6-Hydroxydopamine (6-OHDA) hydrobromide preparation

6-OHDA (Tocris, #2547) was dissolved in a 0.1% (w/v) ascorbic acid (Sigma Aldrich, #AX1775-3) solution in sterile saline to a final concentration of 6 μg/μL. The solution was freshly prepared on the day of surgery, kept on ice, and used within 1.5 hours to maintain stability and potency. Desipramine hydrochloride (Tocris, #3067), a noradrenergic reuptake inhibitor used to prevent damage to noradrenergic terminals, was also prepared (in saline, 10mg/kg dose) and kept on ice.

### Mouse αSyn PFF preparation

Recombinant mouse αSyn PFFs were prepared as described previously^58^. Briefly, monomeric αSyn was overexpressed in *E. coli* BL21 (DE3) cells using the pT7-7 expression vector system and sequentially purified using size exclusion (Cytiva) and HiPrep Q HP 16/10 anion exchange (Cytiva) columns. Endotoxin was removed using endotoxin removal resin (Thermo Fisher Scientific #88277) to a level less than 0.02 units/μg. 0.22μm filter-sterilized αSyn monomers at 5 mg/mL (in phosphate-buffered saline; PBS) were incubated at 37 °C with constant agitation at 1,000 rpm to induce fibrillization. Excess monomer was removed by centrifugation, and fibrils were resuspended in PBS to 5 mg/mL concentration and stored in 25 μL aliquots at –80 °C. Before injection, PFFs were sonicated using a cup horn sonicator (Qsonica #q700) to generate short fibrillar fragments suitable for *in vivo* seeding.

### Stereotaxic surgery

The stereotaxic surgery method for spiny mice was adapted from protocols used for other rodents ^58^. Spiny mice were anesthetized in an isoflurane induction chamber and subsequently transferred to a stereotaxic frame (Kopf Instruments) equipped with a nose cone designed for neonatal rats (RWD #68601). The animals were maintained under isoflurane anesthesia throughout the procedure, with oxygen saturation and heart rate monitored using a MouseSTAT Jr. pulse oximeter and heart rate monitor (Kent Scientific). Isoflurane levels were adjusted to maintain stable anesthesia, with oxygen saturation kept above 95% and heart rate around 350-400 bpm, stabilizing around 250-300 bpm 30-45 minutes after initial induction. Animals were placed on a heating pad to maintain body temperature. Immediately before surgery, ketoprofen (5mg/kg) and buprenorphine-XR (3.25mg/kg) were administered subcutaneously. Desipramine hydrocholoride was administered subcutaneously 30 minutes prior to 6-OHDA injections. The surgical site was disinfected with 3 alternating applications of chlorhexidine scrub and chlorhexidine solutions. Prior to scalp incision, 100 μL of bupivacaine solution (0.25% in sterile saline) was injected subcutaneously over the skull to provide local analgesia and maintain moisture. The skin was incised using surgical scissors rather than a scalpel to minimize shearing, which is a known concern in spiny mouse handling. All surgical steps were performed under sterile conditions, including autoclave sterilization of surgical tools. A 10 μL Hamilton syringe (Hamilton #1701) fitted with a 33-gauge needle was used for stereotaxic injections. Following the injection, the scalp was closed using non-absorbable nylon sutures (Ethicon #NW3353). Before suturing, the animal was gently removed from the ear bars to reduce skin tension and prevent tearing at the incision site. Animals were allowed to recover on a heating pad and monitored for health concerns regularly for the next few days.

The following stereotaxic coordinates and volumes were used for injections: (i) *A. dimidiatus* striatum single-site injection (PFFs; 1.5µL/site): 2.3 AP, LR -2.2 ML, -4.5 DV; (ii) *M. musculus* striatum single-site injection (PFFs; 1.5uL/site): 0.25 AP, -2 ML, -3 DV; (iii) *A. dimidiatus* striatum 2-site injection (6-OHDA :1 μL/site; PFFs:1.5-2 μL/site): 2.31 AP, -2.25 ML, - 4.5 DV & 1.3AP, -3.4 ML, -4.5 DV.

### Open field test

Open field testing was conducted in a square plexiglass arena (22″ x 22″, 12″ height) subdivided to allow simultaneous monitoring of four individual animals. An overhead camera recorded the behavior, and data were analyzed using EthoVision XT 10 (Noldus Information Technology). Animals were acclimated to the behavior room for 30 minutes prior to conducting the behavioral experiment. Each session lasted 12 minutes, with the first 2 minutes designated for habituation within the arena. Locomotor activity and summary statistics were quantified using data from the final 10 minutes of the recording. The arena was cleaned thoroughly between experiments, especially while transitioning between species.

### Hindlimb rearing test

To assess vertical exploratory behavior, animals were placed individually inside a transparent acrylic cylinder (8.5″ diameter, 12″ height) positioned on a clean, stable surface. Each session was recorded using an overhead video camera. Rearing events—defined as the animal lifting both forelimbs off the ground—were counted manually from video recordings. Prior to testing, animals were habituated in the behavior room for 30 minutes and familiarized with the cylinder test setup two days preceding the behavioral test to reduce novelty-induced variability.

### Amphetamine induced rotation

For rotational behavior assessment, animals received an intraperitoneal injection of 2 mg/kg of amphetamine (Sigma-Aldrich#A5880-1G) dissolved in sterile saline. Following the injection, animals were habituated in the behavior room for 30 minutes. Each animal was then placed in an open field arena, and behavior was video recorded using an overhead camera. Rotational events were manually counted over 10 minutes, with rotations defined as full-body turns.

### Immunohistochemistry

At study endpoint, animals were perfused with cold PBS followed by 4% PFA. Brains were post-fixed overnight, cryoprotected in 30% sucrose for 3 days, embedded in OCT, and coronally sectioned at 35 µm using a cryostat. Brain sections stored in PBS with 0.02% azide were permeabilized by incubating the sections in PBS containing 1% Triton X-100 (v/v) (1% PBST) for 1 hour at room temperature. To block non-specific binding, tissues were then incubated in a blocking buffer composed of 10% normal donkey serum (Jackson ImmunoResearch Laboratories #017-000-121) in 0.3% PBST to a primary antibody solution prepared in 0.3% PBST supplemented with 1% normal donkey serum and incubated overnight at 4°C. The next day, sections were washed three times for 10 minutes each in PBS and incubated for 90 minutes at room temperature with fluorophore-conjugated secondary antibodies (Alexa Fluor series, 1:500; Jackson ImmunoResearch Laboratories, West Grove, PA). After secondary labeling, tissues underwent a final series of PBS washes (3 × 10 minutes), were mounted onto glass microscope slides, and left to air-dry overnight. Coverslips were then applied using ProLong™ Diamond Antifade Mountant (Thermo Fisher Scientific, #P36970). Primary antibodies used in this study included: anti-tyrosine hydroxylase (EMD Millipore, #AB152, RRID:AB_390204, 1:1000; #MAB318, RRID:AB_2201528, 1:1000); anti-GFAP (Abcam, #ab4674, RRID:AB_304558, 1:1000), anti-Iba1 (FUJIFILM Wako Pure Chemical Corporation, #019-19741, RRID:AB_839504, 1:1000), anti-pS129 α-Synuclein (EP1536Y) (Abcam #ab51253, RRID:AB_869973, 1:500); anti-pS129 α-Synuclein (81a) (BioLegend #825701, RRID:AB_2564891, 1:500); anti-Ubiquitin (Proteintech #80992-1-RR, RRID:AB_2923694, 1:500); anti-p62/SQSTM1 (Proteintech #18420-1-AP, RRID:AB_10694431, 1:500).

### Imaging

Images were acquired using a spinning disk confocal microscope (Andor, Oxford Instruments) operated with Fusion software (v2.3.0.44) and visualized using Imaris (v9.5.1, Oxford Instruments, RRID: SCR_007370). For striatal density measurements, images were captured using a ChemiDoc imaging system (Bio-Rad) and analysed with ImageJ (NIH, RRID:SCR_002285). The striatal region was defined based on anatomical landmarks, and signal intensity was quantified as mean intensity per unit.

### Laser-capture micro-dissection (LCM) and sample preparation for mass spectrometry

Procedures for tissue excision, lysis, and proteolytic digestion were adapted from previously published work^18^. Briefly, regions of interest from immunohistochemically stained (anti-TH and anti-pS129 αSyn antibodies) tissue sections mounted on PET membranes (Leica Microsystems, cat#76463-322) were isolated using a Leica LMD7 system, collected directly into 50mM TEAB with 0.2% DDM lysis buffer, and stored at -80 °C until processing. For sample preparation, thawed lysates were heat-denatured, enzymatically digested with Lys-C and trypsin, and quenched prior to LC-MS/MS analysis. Comprehensive parameters, including laser settings, buffer composition, incubation conditions, and enzyme amounts, are provided in the protocol reference and in Supplementary Table S1.

### LC-MS/MS analysis

Peptide separations and mass spectrometry analysis were performed as previously described^18^. Briefly, peptides were resolved on an Aurora Ultimate UHPLC C18 column (25 cm × 75 µm, 1.7 µm; IonOpticks #AUR3-25075C18) using a Vanquish Neo UHPLC system coupled to an Orbitrap Astral mass spectrometer (Thermo Scientific) equipped with an EasySpray ion source. Columns were maintained at 55 °C, and 12 µL of each sample was injected for analysis. The LC gradient (Table S2) was operated at a flow rate of 0.20 µL/min for a 19.5-min run. Data-independent acquisition (DIA) was carried out in positive ion mode (1800 V spray voltage, ion transfer tube at 275 °C). MS1 spectra were acquired at 240000 resolution across a scan range of 400-800 m/z with an injection time and an AGC target of 500%. MS2 spectra were acquired on the Astral analyzer with an 80-ms injection time and an AGC target of 800%. Comprehensive DIA parameters and mass spectrometer settings are listed in Table S3.

### Mass spectrometry data analysis and statistics

RAW files acquired in DIA mode were processed with DIA-NN (version 2.2)^59^ using a library-free search with the Swissprot *Mus musculus* database (retrieved 31 July 2025). FASTA digest was enabled with deep learning–based predictions for spectra and retention times. Default DIA-NN parameters were applied with the following modifications: precursor m/z range restricted to 400–800, allowance for up to two missed cleavages. Precursor FDR cutoff was set to 1.0%. Protein and peptide quantification tables were exported for high-confidence proteins (PG.Q.Value < 0.01) for downstream analysis. Only single-cell samples with > 1000 protein IDs were used for analysis, with further stringency applied: (i) isoforms supported exclusively by shared peptides (i.e., without isoform-specific unique peptides) were considered indistinguishable and collapsed as duplicated protein features, and (ii) features (i.e., gene names) were selected if they were present in > 40% of samples in at least one of the groups. MetaboAnalyst (version 6.0, https://www.metaboanalyst.ca, RRID:SCR_015539)^60^ was used for missing value imputation (1/5 Limit-of-Detection method) and median normalization. Statistical analysis and volcano plot generation were performed utilizing NumPy (http://www.numpy.org, RRID:SCR_008633), pandas (https://pandas.pydata.org, RRID:SCR_018214), SciPy (http://www.scipy.org, RRID:SCR_008058), scikit-learn (http://scikit-learn.org, RRID:SCR_002577), seaborn (https://seaborn.pydata.org, RRID:SCR_018132), and scviz packages (https://doi.org/10.5281/zenodo.17362533) in Python (Visual Studio Code [version 1.73, https://code.visualstudio.com, RRID:SCR_026031] or JupyterLab [v3.6.3, http://jupyterlab.github.io/jupyterlab, RRID:SCR_023339] with Python [v3.10.13, http://www.python.org, RRID:SCR_008394] in an Anaconda [v23.11.0, https://docs.anaconda.com, RRID:SCR_025572] environment)^61^. Enrichment analysis was carried out using Kyoto Encyclopedia of Genes and Genomes (KEGG, http://www.kegg.jp, RRID:SCR_012773) pathways as reference. ShinyGO (v0.85, http://bioinformatics.sdstate.edu/go/, RRID:SCR_019213)^62^ was used to perform enrichment and visualize the results as a hierarchical enrichment tree.

### General Statistics

Data were analyzed using unpaired two-tailed Student’s *t*-tests for comparisons between two independent groups, and two-way ANOVA for comparisons involving multiple groups or factors, followed by Tukey’s post hoc analysis where appropriate. Bar graphs were presented as mean ± SEM and box plots indicate the median and interquartile range (IQR; 25th–75th percentile), with the whiskers indicating the minimum and maximum values. The behavioral studies were blinded, and individual animals were not tracked between timepoints (instead considered as a group); therefore, paired analyses were not performed. Statistical analyses were conducted using GraphPad Prism (v9.5.1, http://www.graphpad.com, RRID: SCR_002798) with significance defined as *p* < 0.05.

## Supporting information

Supplementary table S4_Key Resources

Supplementary table S5_protein list

---

**Supplementary Table S1.** Leica LMD7 Settings.

**Laser settings**
|  |  |
| --- | --- |
| **Magnification** | 40x |
| **Power** | 45 |
| **Aperture** | 1 |
| **Speed** | 25 |
| **Specimen balance** | 15 |
| **Line spacing for Draw + Scan** | 19 |
| **Head current** | 100% |
| **Pulse frequency** | 519 |
| **Offset** | 180 |

**Supplementary Table S2.** Gradient conditions for LC-MS/MS analysis on Vanquish Neo-Orbitrap Astral.

**25 min gradient (19.5 min acquisition time)**
| **Time (min)** | **Duration (min)** | **Flow rate (μl/min)** | **% Solvent A** | **% Solvent B** |
| --- | --- | --- | --- | --- |
| **0.0** | 0.0 | 0.450 | 96.0 | 4.0 |
| **0.1** | 0.1 | 0.450 | 96.0 | 4.0 |
| **1.9** | 1.8 | 0.200 | 88.0 | 12.0 |
| **2.0** | 0.1 | 0.200 | 88.0 | 12.0 |
| **13.0** | 11.0 | 0.200 | 77.5 | 22.5 |
| **19.5** | 6.5 | 0.200 | 60.0 | 40.0 |
| **19.5** | **Column Wash** |  |  |  |
| **22.0** | 2.5 | 0.300 | 1.0 | 99.0 |
| **25.0** | 3 | 0.300 | 1.0 | 99.0 |
| **25.0** | **Stop Run** |  |  |  |
| **25.0** | **Column Equilibration** |  |  |  |

**Supplementary Table S3.** MS settings for DDA and DIA LC-MS/MS analysis on Vanquish Neo-Orbitrap Astral.

| Settings | Full scan | DIA (MS2) |
| --- | --- | --- |
| Resolution | 240000 |  |
| AGC target (%) | 500% | 800% |
| Maximum injection time | 100 ms | 80 ms |
| Scan range | 400-800 m/z | 150-2000 m/z |
| Isolation window (m/z) |  | 20 m/z |
| Precursor mass range |  | 400-800 m/z |
| RF lens (%) | 45 | 45 |
| HCD collision energy (%) |  | 25 |

**Supplementary Figure 1.**
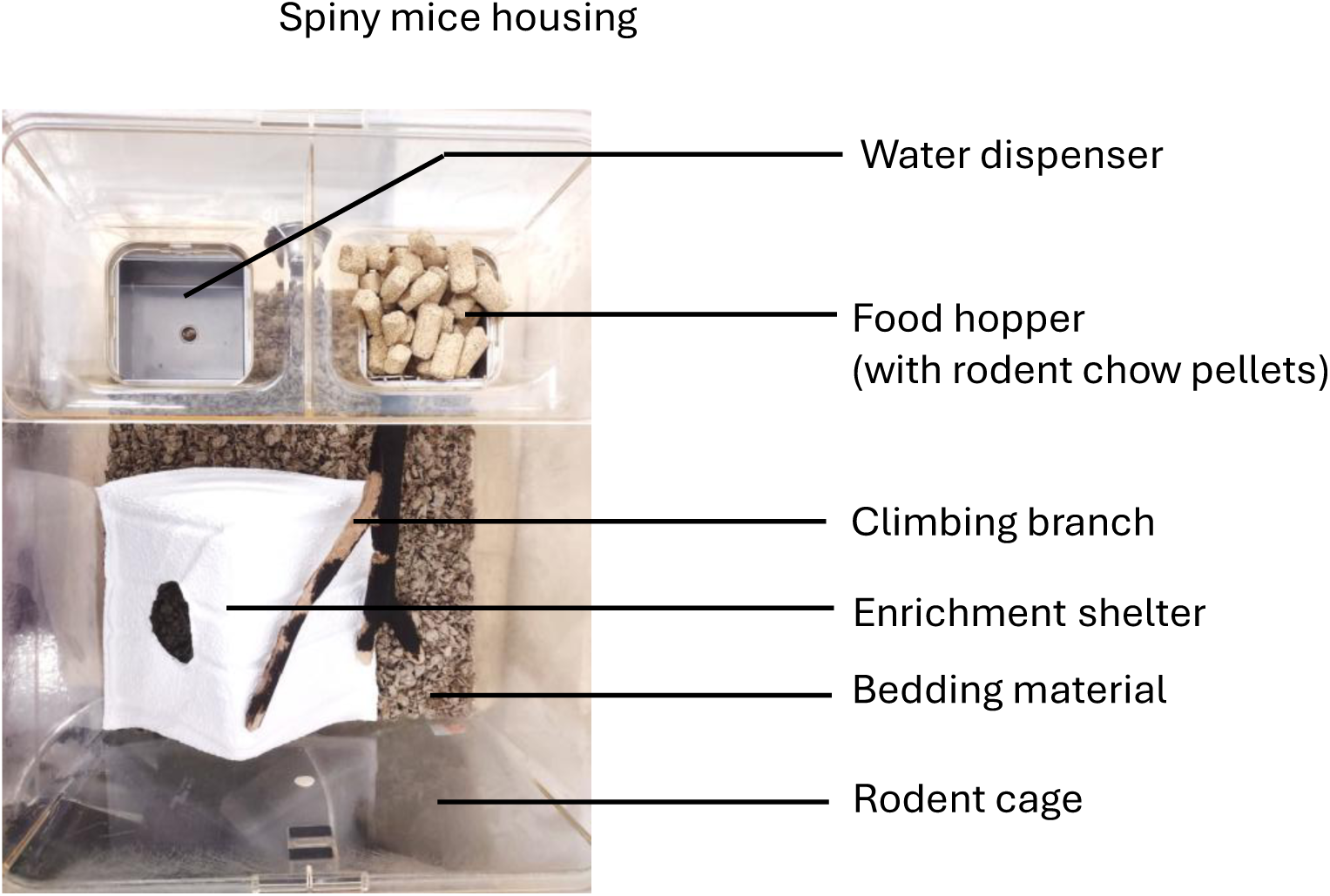
Small-scale group housing setup for adult spiny mice (*Acomys dimidiatus*). Photograph showing a representative housing configuration for 3–5 adult spiny mice in a standard rat cage (Lab Products “Super Rat” polycarbonate cages, dimensions: 13×11×8”). The cage is equipped with a feeder for dry chow and a water pouch for hydration for breeding cages and autoclaved water for non-breeding cages. Environmental enrichment includes a wooden block, a manzanita branch to encourage climbing behavior, and a Shepherd Shack shelter to support nesting and reduce stress. This housing arrangement supports the social and exploratory behavior typical of *Acomys*.

**Supplementary Figure 2.**
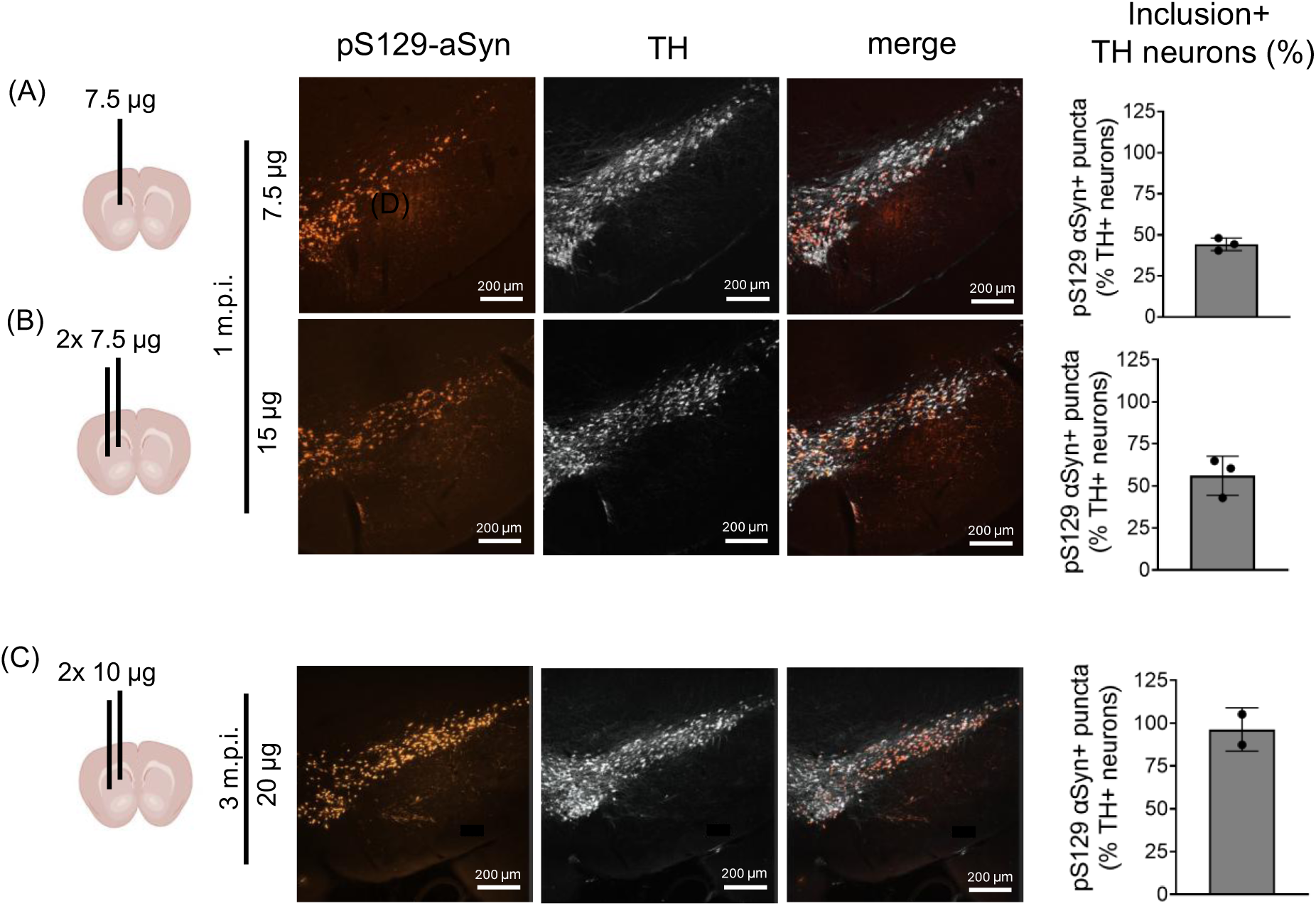
Dose-dependent induction of pS129-αSyn pathology in *A. dimidiatus* following striatal αSyn PFF injection. Schematic diagram illustrates the injection locations for various doses of αSyn PFFs delivered to the striatum. Representative coronal brain sections are shown for each condition, depicting pS129-αSyn aggregation (orange) alongside TH+ cell bodies (white). Quantification of pS129-αSyn+ cell bodies is presented as a percentage of total TH⁺ neurons in the SNpc for the following conditions: (A) 7.5 µg PFFs (single-site injection, 1-month post-injection); (B) 15 µg PFFs (two-site injection, 1-month post-injection); (C) 20 µg PFFs (two-site injection, 3 months post-injection).

**Supplementary Figure 3:**
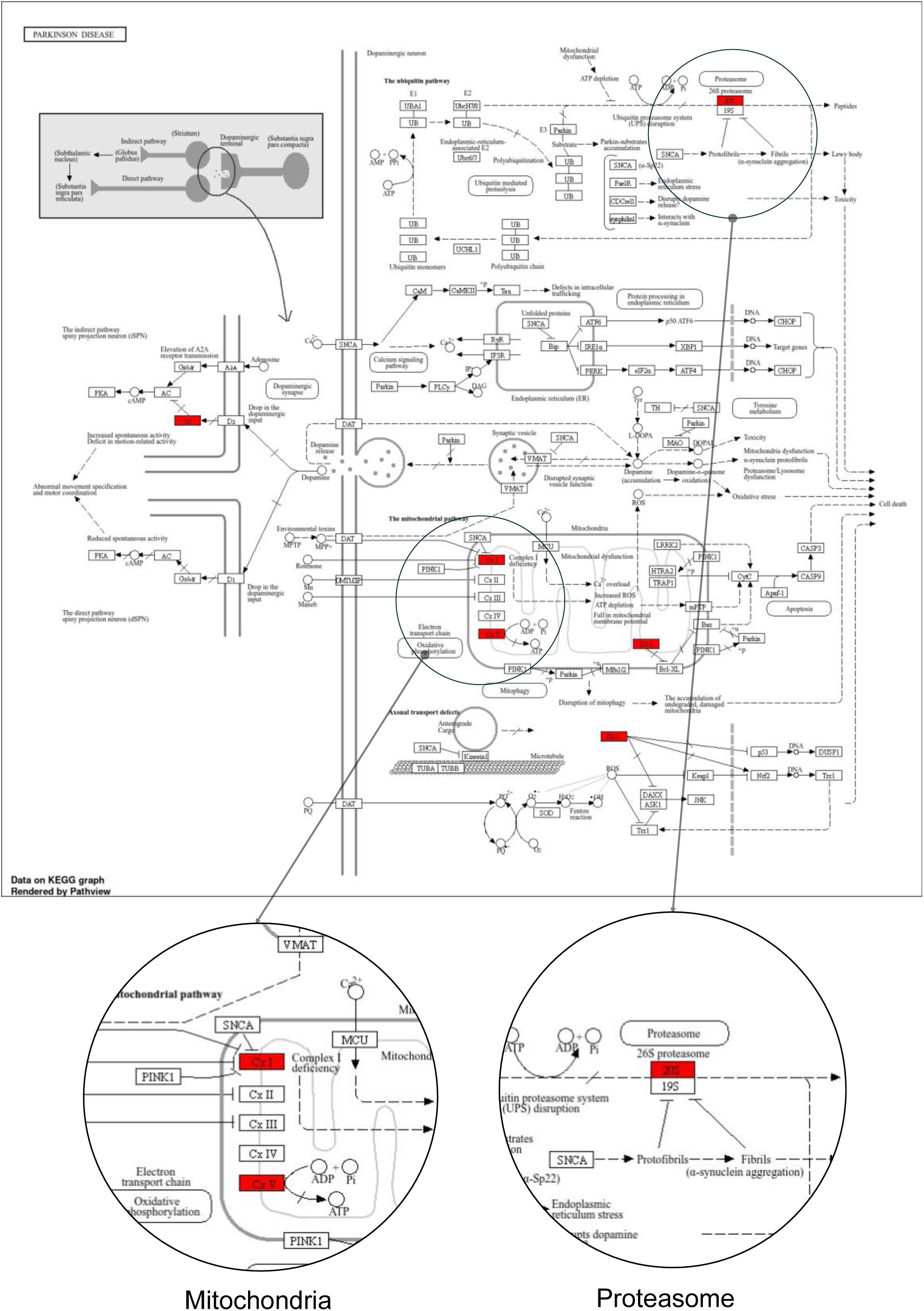
KEGG Parkinson’s disease (has05012) pathway. Red-highlighted boxes indicate proteins significantly altered by αSyn aggregates in *A. dimidiatus* SNpc (p ≤ 0.05, fold change ≥ 2) within the KEGG “Parkinson’s disease” pathway.

**Supplementary Figure 4:**
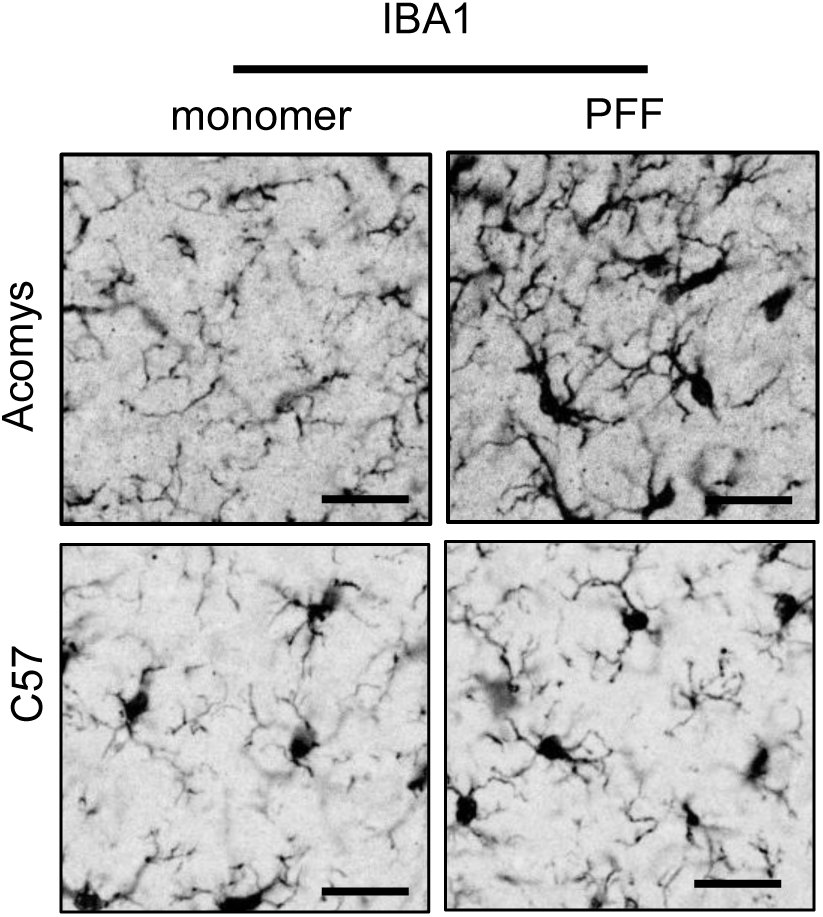
Microglial immunoreactivity. Representative microglial IBA1 immunostaining in the substantia nigra of spiny mice and C57BL/6J mice at 6 months post-PFF injection. Images show the distribution and relative density of IBA1-positive microglia within the substantia nigra. Scale bars, 25 µm.

## Acknowledgments

This research was funded in whole by Aligning Science Across Parkinson’s (ASAP-020495 to V.G.) through the Michael J. Fox Foundation for Parkinson’s Research (MJFF). M.P. was supported by the A*STAR National Science Scholarship (BS-PhD). For the purpose of open access, the authors have applied a CC BY public copyright license to all Author Accepted Manuscripts arising from this submission.

We thank Leica Biosciences for access to the LMD7 instrument. We thank Caltech’s Office of Laboratory Animal Research (OLAR) for support in maintaining the *A. dimidiatus* colony at Caltech. We thank Caltech’s Proteome Exploration Laboratory (PEL) for support and access to mass spectrometry instruments.

## Author contributions

S.D. conceived the overall study, designed the experiments, performed animal husbandry, conducted stereotaxic surgery, prepared α-synuclein preformed fibrils, conducted behavioral experiments, tissue processing, immunohistology staining, and imaging, and prepared the manuscript with input from all authors. S.D. and M.P. performed LCM sample collection, data analysis, and visualization. M.P. conducted all MS raw data processing. R.R.D. and A.W.S. advised on spiny mouse model and colony maintenance, behavior, and experimental design. T-F.C. advised on proteomics experiments and data analysis. V.G. supervised the overall study and advised on experimental design. All authors reviewed and edited the manuscript.

## Data Availability

The data, protocols, and key lab materials used and generated in this study are listed in a Key Resource Table alongside their persistent identifiers in Supplementary Table S4. A complete list of identified proteins, including differentially expressed proteins with their fold changes and p-values for *A. dimidiatus*, is provided in Supplementary Table S5

